# CRISPR Mutants of Three Y Chromosome Genes Suggest Gradual Evolution of Fertility Functions in *Drosophila melanogaster*

**DOI:** 10.1101/2020.01.30.926543

**Authors:** Yassi Hafezi, Samantha R. Sruba, Steven R. Tarrash, Mariana F. Wolfner, Andrew G. Clark

## Abstract

Gene-poor, repeat-rich regions of the genome are poorly understood and have been understudied due to technical challenges and the misconception that they are degenerating “junk”. Yet multiple lines of evidence indicate these regions may be an important source of variation that could drive adaptation and species divergence, particularly through regulation of fertility. The ∼40 Mb Y chromosome of *Drosophila melanogaster* contains only 16 known protein-coding genes and is highly repetitive and entirely heterochromatic. Most of the genes originated from duplication of autosomal genes and have reduced nonsynonymous substitution rates, suggesting functional constraint. We devised a genetic strategy for recovering and retaining stocks with sterile Y-linked mutations and combined it with CRISPR to create mutants with deletions that disrupt three Y-linked genes. Two genes, *PRY* and *FDY*, had no previously identified functions. We found that *PRY* mutant males are sub-fertile, but *FDY* mutant males had no detectable fertility defects. *FDY*, the newest known gene on the Y chromosome, may have fertility effects that are conditional or too subtle to detect. The third gene, *CCY*, had been predicted but never formally shown to be required for male fertility. CRISPR-targeting and RNAi of *CCY* caused male sterility. Surprisingly, however, our *CCY* mutants were sterile even in the presence of an extra wild-type Y chromosome, suggesting that perturbation of the Y chromosome can lead to dominant sterility. Our approach provides an important step toward understanding the complex functions of the Y chromosome and parsing which functions are accomplished by genes versus repeat elements.

## INTRODUCTION

Y chromosomes are rapidly evolving and may contribute to the diversification and adaptation of species, yet functional characterization of these regions lags far behind the rest of the genome (Hughes *et al*. 2010; Bachtrog 2013; Hughes and Page 2015; Tobler *et al*. 2017). In *Drosophila melanogaster* the ∼40 Mb Y chromosome is entirely composed of constitutive heterochromatin (Heitz 1933), densely populated by repetitive sequences (Lohe *et al*. 1993; Hoskins *et al*. 2015; Chang and Larracuente 2019), and contains only 16 known protein-coding genes (Gepner and Hays 1993; Carvalho *et al*. 2000, 2001, 2015; Vibranovski *et al*. 2008; Krsticevic *et al*. 2010). This scarcity is caused by the highly unusual genetic environment where there is no recombination, male-restricted selection, and one-fourth of the autosomal effective population size. Both classic genetic and genomic approaches have been used to dissect the function and composition of the Y chromosome, respectively. Despite these efforts, very little is understood about how specific genetic elements fulfill the functions ascribed to the Y.

Collectively, the Y chromosome fulfills several important functions. The Y is essential for male fertility (Bridges 1916a; b), due to several genetic loci, called “fertility factors” (Brosseau 1960; Kennison 1981). “Saturation” or the existence of multiple alleles of each fertility factor (including X-ray, P-element, EMS, and segmental deletion) strongly suggests there are only six fertility factors on the Y (Ayles *et al*. 1973; Hazelrigg *et al*. 1982; Kennison 1983; Gatti and Pimpinelli 1983; Zhang and Stankiewicz 1998). Variation on the Y also contributes quantitatively to male fertility and fitness (Clark 1990; Chippindale and Rice 2001), temperature adaptation of spermatogenesis (Rohmer *et al*. 2004), global and testis-specific gene expression regulation (Zhang *et al*. 2000; Lemos *et al*. 2008, 2010), position effect variegation/chromatin regulation (Dimitri and Pisano 1989, Kelsey and Clark, in prep), geotaxis/locomotor activity (Stoltenberg and Hirsch 1997; Dean *et al*. 2015) and sex-specific aging (Griffin *et al*. 2015; Brown and Bachtrog 2017).

There are several challenges in studying the Y chromosome. Repetitive AT-rich sequences complicate PCR amplification, sequencing and assembly (Hoskins *et al*. 2015). Heterochromatic condensation of the Y obscures visible markers and prevents endoreduplication (Smith and Orr-Weaver 1991; Belyaeva *et al*. 1998), thereby reducing sequence representation. Recently, long-read sequencing was used to improve the coverage and contiguity of the Y chromosome sequence by over three fold (Chang and Larracuente 2019). Still only ∼14.6 out of 40 Mb of Y chromosome sequence are in the latest assembly. Hemizygosity complicates the recovery, maintenance and complementation analysis of mutants. And traditional genetic mapping is not possible in the absence of recombination (Kennison 1981).

The identification of protein-coding sequences on the Y chromosome was a major breakthrough involving multiple computational strategies, i.e. analyzing contrasts between male and female DNA and RNA sequences (Carvalho and Clark 2013). The 13 identified single-copy genes fall into two categories: six are predicted to be fertility factors and the remaining seven genes have unknown function. The genes identified on the Y chromosome from sequencing were mapped against segmental deletions, large deletions created by intercrossing fertile X-Y chromosome translocations (Kennison 1981; Hardy *et al*. 1981) (Table S1). These regions are large, some contain multiple genes, and with only ∼36.5% of the Y chromosome assembled additional genes are expected to be discovered (Chang and Larracuente 2019). Moreover, exciting new findings suggest that RNA transcripts from some simple tandem repeats can be required for male fertility in *Drosophila melanogaster* (Mills *et al*. 2019) challenging the assumption that the fertility factors must be protein-coding genes.

Thus, although candidate genes corresponding to each fertility factor have been identified (Table S1), functional evidence is needed to definitively link individual genes to the sterility phenotype. Functional evidence suggests that two dynein genes, *kl-3* and *kl-5*, are fertility factors (Ayles *et al*. 1973; Goldstein *et al*. 1982; Gepner and Hays 1993; Yu *et al*. 2013). Four other genes, *WDY, kl-2, ORY*, and *CCY*, were predicted to be fertility factors, corresponding to *kl-1, kl-2, ks-1*, and *ks-2*, respectively (Carvalho *et al*. 2000, 2001; Vibranovski *et al*. 2008), but have not been functionally tested.

The remaining seven genes are likely not required for the production of offspring – *PRY*, for example, is located at the breakpoint of a translocation line that is fertile (Carvalho *et al*. 2000). Yet these genes are still expected to contribute, albeit more subtly, to male fertility and/or fitness. This is due to the male-limited inheritance, testis-biased expression (Brown *et al*. 2014; Mahajan and Bachtrog 2017) and evidence of selection on all Y genes. Reduced nonsynonymous substitution rates of Y-linked orthologs have been reported for *PRY, ARY*, and *Ppr-Y* (Singh *et al*. 2014). *FDY* is only found in *Drosophila melanogaster*, but the preservation of its large open reading frame is statistically implausible under neutrality and is therefore evidence for functional constraint (Carvalho *et al*. 2015). Finally, the majority of the genes on the Y chromosome were acquired through duplications or retrotranspositions from autosomes, most recently within the past 2.5 million years (Koerich *et al*. 2008; Tobler *et al*. 2017).

Here we investigated the role of specific Y chromosome genes in male fertility by using CRISPR to disrupt the genes. CRISPR has been shown to work on the Y chromosome for inducing double-strand breaks followed by nonhomologous end joining (Yu *et al*. 2013) and for site-specific transgene insertion, albeit relatively inefficiently (Buchman and Akbari 2018). We target three genes – *CCY, PRY* and *FDY* – which span a range of ages of Y-linkage (Koerich *et al*. 2008) (Table 1). *FDY* is the youngest known Y chromosome gene (Carvalho *et al*. 2015). In contrast, *PRY* was present on the ancestral Y chromosome, though it has been lost in some species (Koerich *et al*. 2008). *CCY* was acquired after the Sophophora split 63 million years ago and is unique in having no obvious autosomal paralog in *Drosophila melanogaster*. We identify strong fertility defects in mutants of the two older genes and no detectable fertility defect in mutants of the youngest gene, which suggests there may be a gradual gain of fertility functions in the evolution of Y chromosome genes.

**Table 1.**
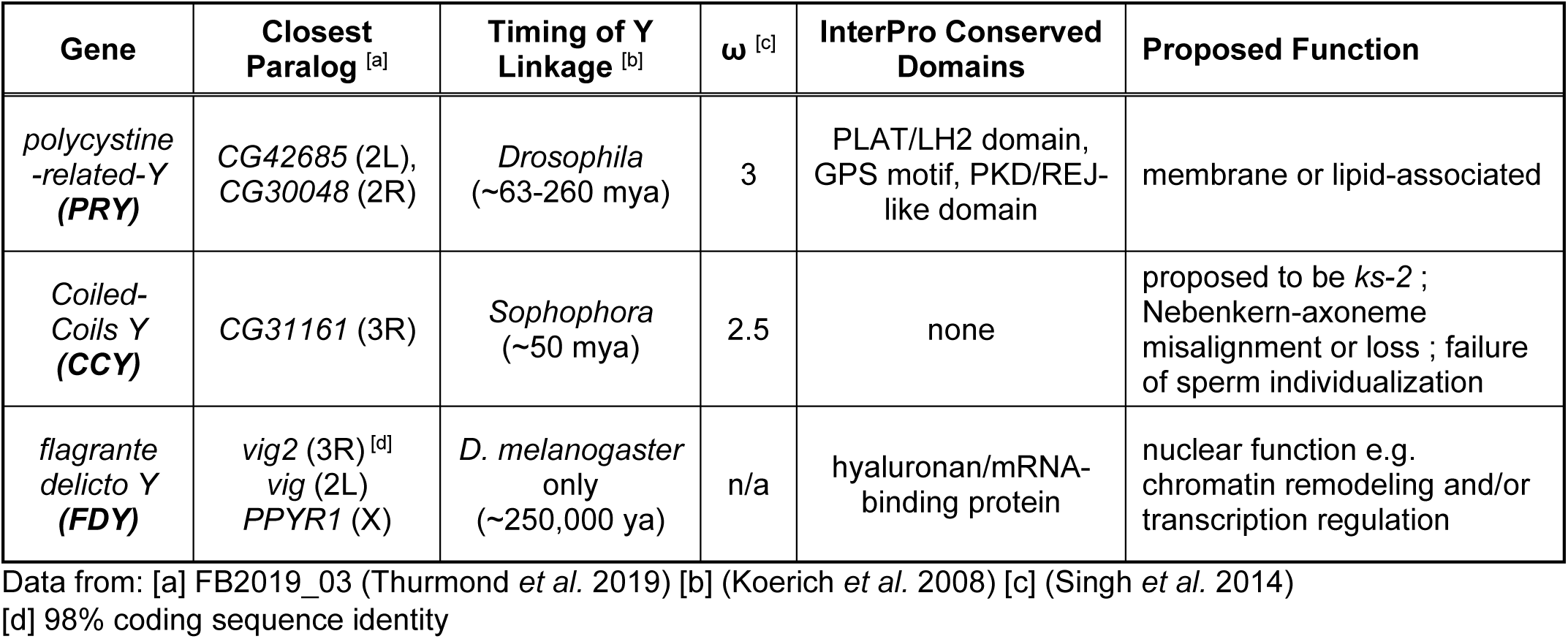
Summary of the genes investigated and their proposed functions.

| Gene | Closest Paralog <sup>[a]</sup> | Timing of Y Linkage <sup>[b]</sup> | $\omega$ <sup>[c]</sup> | InterPro Conserved Domains | Proposed Function |
| --- | --- | --- | --- | --- | --- |
| *polycystine-related-Y (PRY)* | CG42685 (2L),<br>CG30048 (2R) | *Drosophila*<br>(~63-260 mya) | 3 | PLAT/LH2 domain,<br>GPS motif, PKD/REJ-like domain | membrane or lipid-associated |
| *Coiled-Coils Y (CCY)* | CG31161 (3R) | *Sophophora*<br>(~50 mya) | 2.5 | none | proposed to be *ks-2* ;<br>Nebenkern-axoneme<br>misalignment or loss ; failure<br>of sperm individualization |
| *flagrante delicto Y (FDY)* | *vig2* (3R) <sup>[d]</sup><br>*vig* (2L)<br>*PPYR1* (X) | *D. melanogaster*<br>only<br>(~250,000 ya) | n/a | hyaluronan/mRNA-binding protein | nuclear function e.g.<br>chromatin remodeling and/or<br>transcription regulation |

Data from: [a] FB2019\_03 (Thurmond *et al.* 2019) [b] (Koerich *et al.* 2008) [c] (Singh *et al.* 2014)
[d] 98% coding sequence identity

## MATERIALS AND METHODS

### Fly stocks and husbandry

Flies were reared on a cornmeal-agar-sucrose medium (recipe available at https://cornellfly.wordpress.com/s-food/) at 25°C with a 12 h light-dark cycle. The RNAi lines used are listed in Table 4, and the remaining fly lines used are in Table S2.

**Table 2.** Summary of the *PRY* and *FDY* alleles generated by CRISPR. No alleles of either gene resulted in sterile males.

| Allele | Bases Deleted | Bases Inserted | Total Bases Change | Effect | Sterility Test Result |  |  |
| --- | --- | --- | --- | --- | --- | --- | --- |
|  |  |  |  |  | # | % |  |
| *PRY*Control | - | - | - | - | 20/20 | 100% | **Not Sterile** |
| *PRY* G5.7 | 241 | 0 | -241 | Out-of-frame | 10/10 | 100% | **Not Sterile** |
| *PRY* G54.1 | 244 | 12 | -232 | Out-of-frame | 9/10 | 90% | **Not Sterile** |
| *PRY* G54.7 | 259 | 1 | -258 | In-frame | 6/10 | 60% | **Not Sterile** |
| *PRY* G55.8 | 240 | 1 | -239 | Out-of-frame | 10/10 | 100% | **Not Sterile** |
| *PRY* G61.2 | 239 | 9 | -230 | Out-of-frame | 10/10 | 100% | **Not Sterile** |
| *FDY*Control | - | - | - | - | 25/25 | 100% | **Not Sterile** |
| *FDY* L3.2 <sup>[a]</sup> | 587 | 0 | -587/-382 <sup>[b]</sup> | Out-of-frame | 75/75 | 100% | **Not Sterile** |
| *FDY* L59.1 | 579 | 0 | -579/-375 <sup>[b]</sup> | In-frame | 10/10 | 100% | **Not Sterile** |
| *FDY* L64.3 | 581 | 2 | -579/-375 <sup>[b]</sup> | In-frame | 5/5 | 100% | **Not Sterile** |
| *FDY* L71.4 | 595 | 2 | -593/-383 <sup>[b]</sup> | Out-of-frame | 5/5 | 100% | **Not Sterile** |

[a] Six independently-derived lines had lesions identical to *FDY* L3.2.
[b] Second number represents the total BP change from the start of the coding sequence to the 3' break-point. This number was used for determining if a frame shift occurred.

**Table 3.** Mutations that disrupt *CCY* reading frame result in male sterility despite the presence of a wild-type copy of *CCY*.

|  | **Total Flies Sequenced** | **Flies With In-Frame INDELS** | **Flies With Out-Of-Frame INDELS** | **Flies With No Double Peaks <sup>[a]</sup>** |
| --- | --- | --- | --- | --- |
| Not sterile | 14 | 9 | 0 | 5 |
| Sterile | 16 | 0 | 9 | 7 |
| **Total** | **30** | **9** | **9** | **12** |

[a] INDELS on the free Y in $\widehat{X}YY$ males resulted in double peaks on sequence traces. A subset of flies, both sterile and non-sterile, had no double peaks and therefore no recognizable lesion.

**Table 4.**
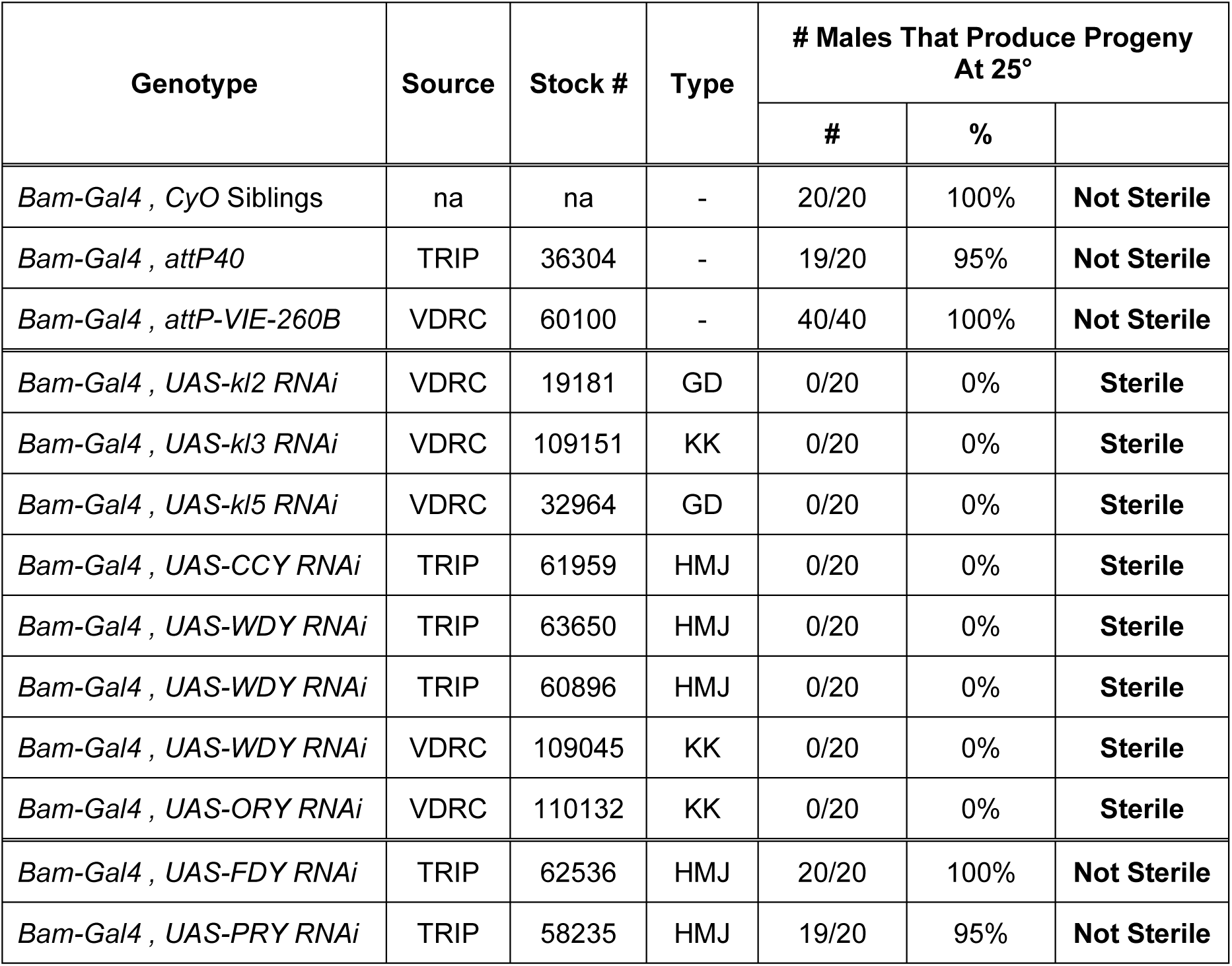
RNAi of *CCY* and the other predicted fertility factors sterilizes males while RNAi of *PRY* and *FDY* does not.

The “balanced double compound” (BDC) line (Fig. 1) was constructed with C(1)M4 (referred to as 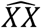 and C(1;Y)1 (referred to as 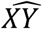) chromosomes. To create this line we first crossed Canton-S males to females from each compound chromosome line (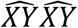 and 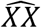) to recover flies with an extra Canton-S-derived Y chromosome (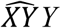 and 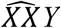). We then crossed the 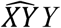 males and 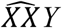 females together to establish the BDC line. We only used BDC lines constructed within 6 months of injections to avoid the accumulation of Y-linked mutations that might occur when an extra Y chromosome is present.

**Figure 1.**
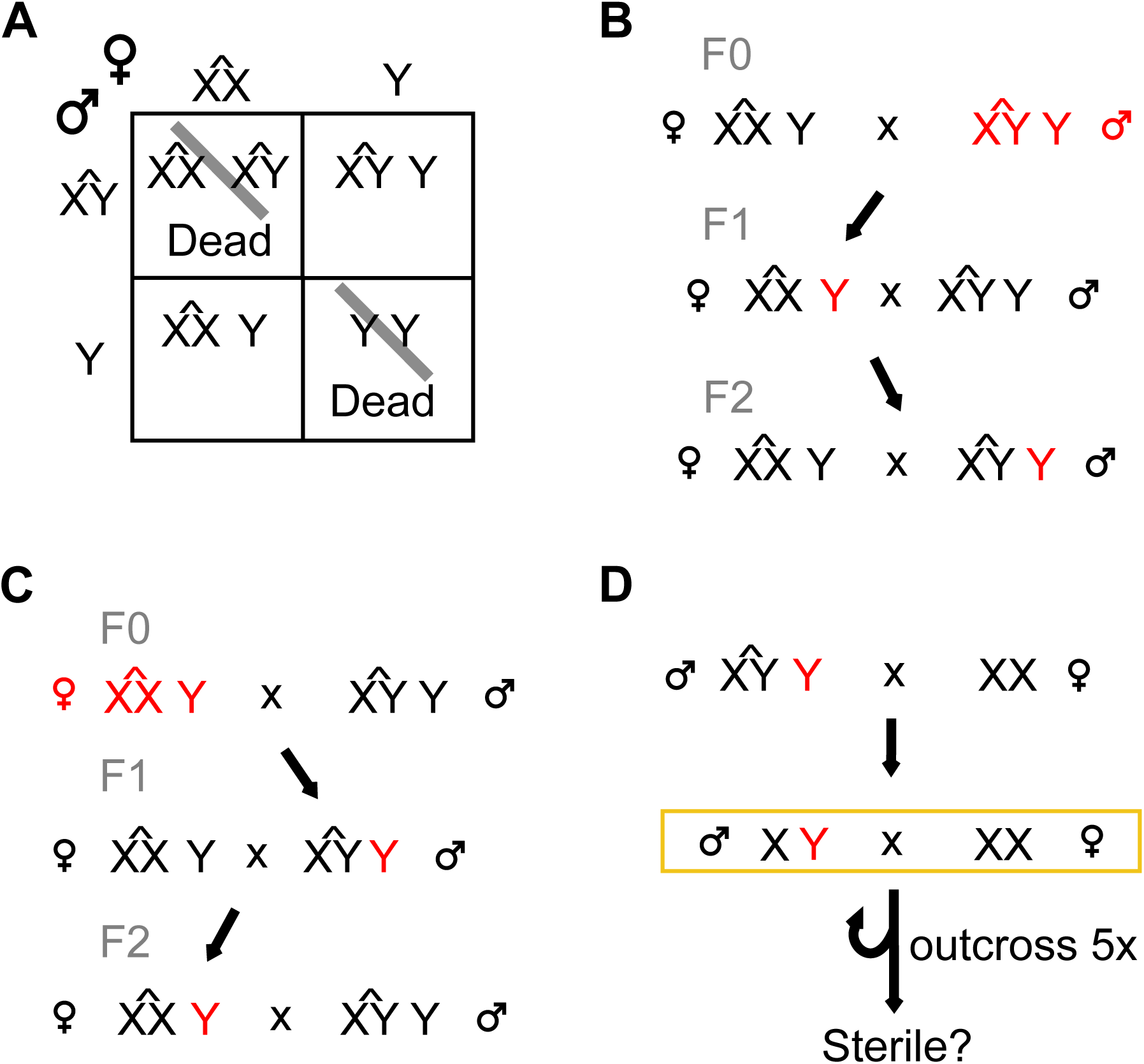
Crossing scheme for CRISPR targeting strategy using the BDC line. (A) In the BDC line an extra, free Y chromosome is maintained stably and passed to the opposite sex at each generation. (B,C) Editing occurs in flies injected with Cas9 and guide RNAs (red). (B) Males have two Y chromosomes that can be edited, while (C) females have a single non-essential Y chromosome. (D) Once a stable mutant is confirmed, the Y is removed from the compound background, tested for sterility (yellow box), and outcrossed to Canton-S for a minimum of 5 generations.

Images of adult eyes were captured with a Canon Rebel 6-megapixel digital camera attached to an Olympus SMZ-10 dissecting microscope.

### CRISPR Mutants

We targeted roughly 1 kb regions at or near the 5’ end of the coding sequence of each gene. Each target region was first amplified in males and females to confirm male-specificity and sequenced to identify any polymorphisms in the Canton-S strain and fill in any sequence gaps. Single guide RNAs (sgRNAs) were designed using ‘CRISPR Optimal Target Finder’ (http://tools.flycrispr.molbio.wisc.edu/targetFinder/). We aimed to design sgRNAs that were 20 nt long, contained at least one 5’ G, and had no off-target sites in the Drosophila genome. When possible, we opted for 4 G/Cs in 6 bases adjacent to the PAM site (Ren, 2014). If the sgRNA only had one 5’ G, a second G was added to create an optimal T7 promoter. Our primer and sgRNA sequences are listed in Table S3.

We aimed for unique sgRNA sequences, especially in the nucleotides adjacent to the PAM motif, as these are more important for target recognition (Ren 2014). However, due to the sequence similarity of *FDY* and *vig2*, we could not find an sgRNA that targeted one gene but not the other. We therefore used sgRNAs that would target both. After six outcrosses (an extra outcross was performed for these mutants) we confirmed that our *FDY* mutants did not contain any mutations in the homologous *vig2* locus.

sgRNAs were synthesized as described in Kistler *et al*. (2015). Briefly, a DNA template was made by template-free PCR and used in an in vitro transcription reaction (T7 MEGAscript kit, LifeTechnologies AM1334). The RNA was purified on a column (MEGAclear column, LifeTechnologies AM1908) and by sodium acetate precipitation. An injection mix was made with 40 ng/ul sgRNA RNA and 300 ng/ul Cas9 protein (PNA Bio # CP01-50) and injected into embryos from our BDC stock (Fig. 1) by Rainbow Transgenic Flies, Inc.

Single male and female individuals from the F0 (injected) generation were crossed to the BDC parental line as show in Fig. 1B and C in order to isolate and propagate the edited Y chromosome. The individual flies from the F0 and F1 generations were genotyped by PCR once larvae of the next generation were apparent.

Generally, uninjected BDC flies were used as controls (e.g. *PRY* control), however, for the *FDY* mutants, we used flies that were injected with Cas9 and sgRNAs but not edited at the *FDY or vig2* locus as our controls. The extra Y chromosome was removed from controls and control flies were outcrossed alongside the respective mutant.

### Fertility Assays

To test if a particular genotype (RNAi-expressing or mutant) was sterile, we crossed 15-20 males at three to five days old to five Canton-S virgin females in a vial for one week, then transferred the adults into a new vial for another week. If no progeny were observed in either vial the male was considered sterile. No quantitative assessment was made at this point. Vials with no progeny whose male died within the test period were not counted.

For a more quantitative measure of fertility (CRISPR mutants only) we mated three mutant or wild-type males, three to five days old, to single three to four day old Canton-S virgin females. Flies were observed for several hours; females that mated were kept while, females that did not mate and all males were discarded. Females were transferred to a new vial every day for five days and discarded on the fifth day after mating. Fresh active yeast mixed 1:1 with water was provided each day, starting two days before mating. The number of eggs laid (fecundity) and pupae emerging (fertility) from each vial was counted. Hatchability was calculated for each group as mean number of pupae divided by the mean number of eggs laid. We tested three different alleles of each genotype with two independent replicates of each allele. We first confirmed by a Kruskal-Wallis rank sum test that the independent alleles of each genotype were not statistically different. A Wilcoxon rank sum test was then applied to compare total eggs laid and pupae produced over the five days between the mutant and control groups. To detect differences in the variability of fertility or fecundity of the mutant and control groups, we used the Levene Test for homogeneity of variance.

### Transcript Levels

To estimate transcript levels we first extracted RNA from 20-25 adults or pairs of testes using TRIzol. The RNA was treated with DNase (Promega #M6101, RQ1 RNase-Free DNase) and used to synthesize cDNA (Takara #639537, SMARTScribe Reverse Transcriptase). PCR was used to amplify gene-specific sequences and results were analyzed on a 1.5% agarose gel. Transcript levels were estimated by comparing the band intensity of undiluted mutant or RNAi samples to that of a dilution-series from the corresponding control. We controlled for contaminating genomic DNA in each sample by including a cDNA synthesis reaction without reverse transcriptase. Housekeeping gene transcripts (*Actin5C* or *RpL32*) were used as standards to test for even loading of cDNA between samples. All primer sequences used are provided in Table S3.

### Cytology

Fixed spermatocyte squash preparations were made as described in Sitaram *et al*. (Sitaram *et al*. 2014). Briefly, testes were dissected from adult males within 24 h of eclosion. They were torn half-way between the coil and anterior tip and squashed under a coverslip. Slides were snap frozen and coverslips were removed. Slides were then immersed in cold ethanol, fixed for 7 min in 4% paraformaldehyde in PBS, washed in 0.1% Triton-X in PBS, and blocked with 1% BSA. Spermatocytes were stained with 0.01 ug/mL phalloidin-TRITC (Sigma-Aldrich, P1951), mounted in Vectashield with DAPI and imaged on a Zeiss 710 confocal microscope. We karyotyped mitotic chromosomes by using the method in Bauerly *et al*. (Bauerly *et al*. 2014). Briefly, we separated male and female wandering third instar larvae and dissected brain complexes in PBS. Brains were incubated in colchicine for 1 h, 0.5% sodium citrate for 6 min, and fixed in 45% acetic acid + 3% paraformaldehyde for 5 min. Brains were then squashed, frozen in liquid nitrogen, mounted in Vectashield with DAPI and imaged on an Olympus fluorescence microscope.

### Data availability

The authors affirm that all data necessary for confirming the conclusions presented in the article are represented fully within the article and supplemental material. Supplemental Figures S1-S6 and Supplemental Tables S1-S10 are available through FigShare. The reported mutants are available upon request.

## RESULTS

### CRISPR strategy for making Y chromosome gene mutants

We expected fertility factor genes to be recessive sterile and mutations in any other Y chromosome genes to potentially reduce male fertility (Carvalho *et al*. 2001). To allow recovery of such mutants we needed to create them in either females or in males with an extra Y chromosome. We created a “balanced double compound” (BDC) line where males and females each have a compound sex chromosome (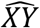 and 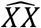, respectively) as well as an extra free Canton-S-derived Y chromosome (Fig. 1A). Depending on the efficiency of Cas9 cleavage, neither, one or both copies of the Y chromosomes in F0 males might be mutated. Mutation of both Y chromosomes in males may lead to their sterility, but we can retain all Y chromosome mutations arising in F0 female flies. Lesions inherited from F0 females will be passed to F1 males that contain an additional wild-type Y chromosome inherited from the 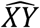 of unperturbed BDC fathers (Fig. 1C). The compound 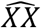 chromosome additionally has a *w*^*m4*^ allele to allow for visual screening for loss of the Y chromosome, which would result in a loss of suppression of position effect variegation by the Y. Thus, 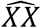 females have mostly-white eyes while 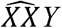 females have mostly-red eyes (Fig. 3A,B). Both males and females from the BDC line were edited and mutants were recovered (Fig. 1B,C).

**Figure 2.**
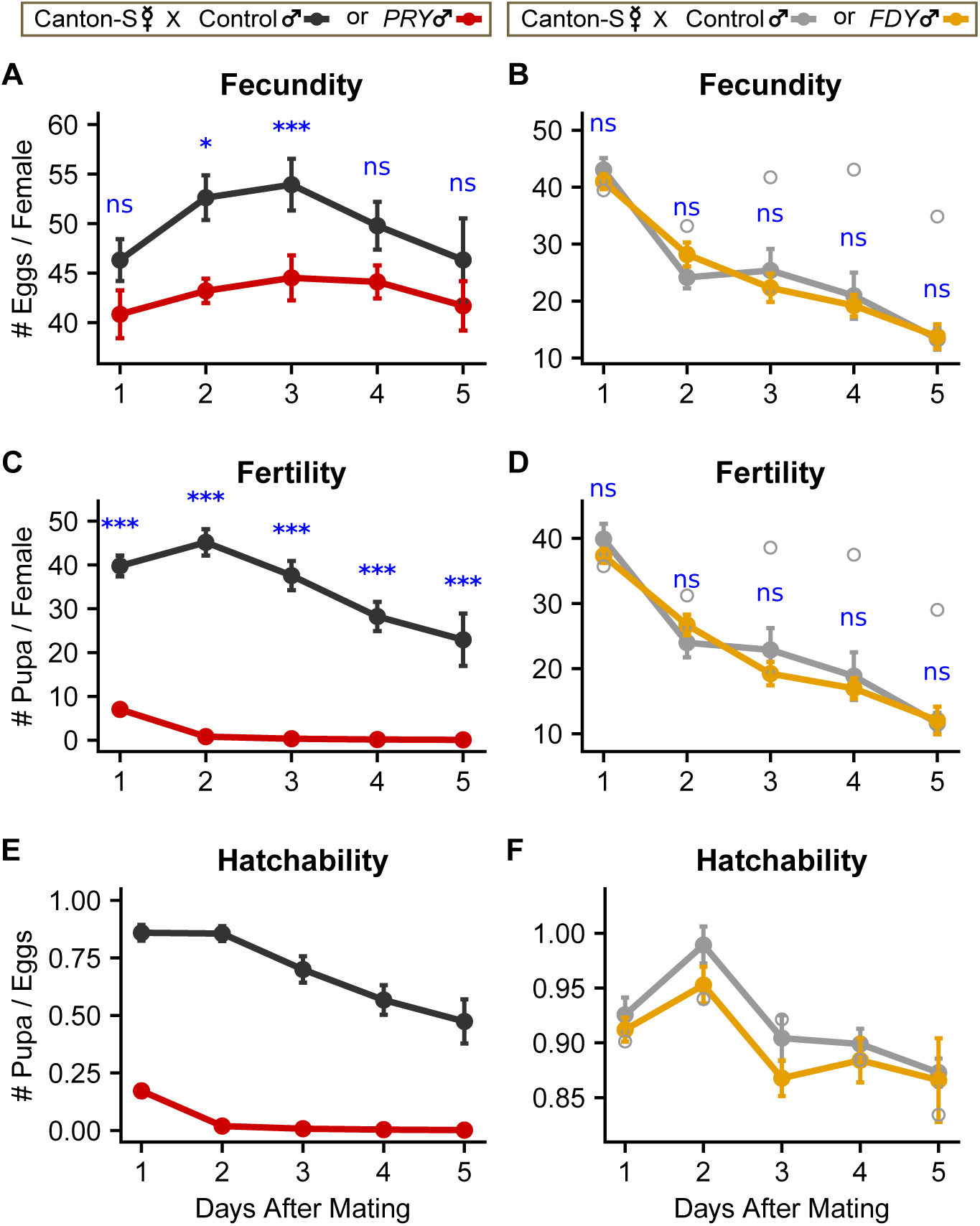
*PRY* activity contributes to male fertility, while *FDY* activity has no detectable effect on male fertility. (A,B) Fecundity (number of eggs laid), (C,D) fertility (number of pupae produced), and (E,F) hatchability (fertility/fecundity) of *PRY* (A,C,E) or *FDY* (B,D,F) compared to control males crossed to Canton-S virgin females. Closed circles represent the overall mean for all alleles and repeats per genotype per day. Error bars represent standard error of the mean. In some cases, error bars are not visible because they are smaller than the symbol size for the mean. Open circles represent the mean of an outlier allele, *FDY’s* control allele E, that was omitted. Results of the Wilcoxon Rank Sum Test for significant differences between genotypes are indicated in blue. * p<0.05, *** p<0.01, ns = not significant.

**Figure 3.**
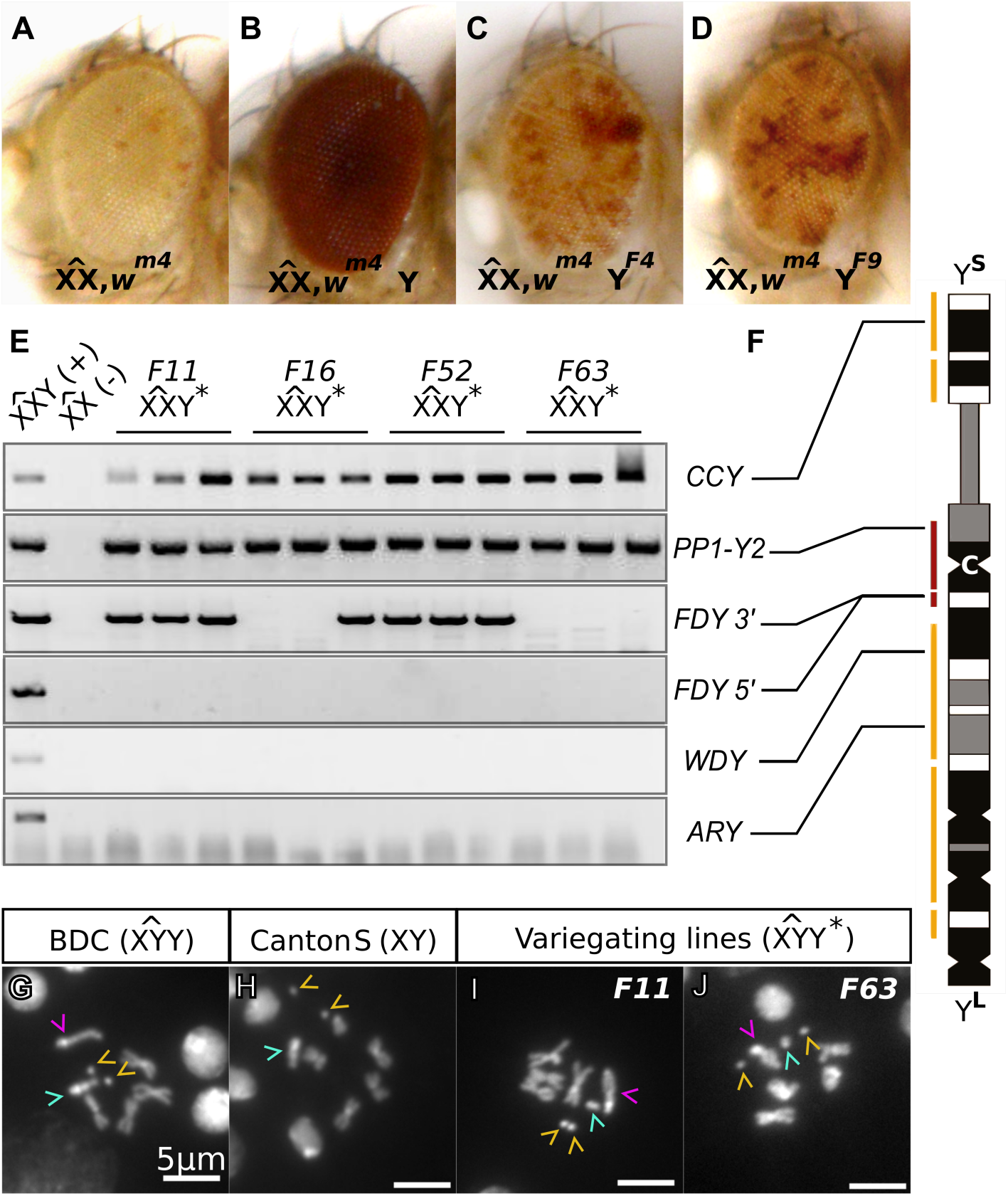
Variegation artifact during creation of *FDY* mutants caused by Y chromosome truncation. (A-D) Adult female eyes of controls (A,B) and two different variegating lines (C,D) that arose when we targeted *FDY*. (E) PCR amplification of three variegating females each from four independently-derived variegating lines. (F) Map of the scaffolds (yellow) and contigs (red) in the most recent sequence assembly of the Y chromosome (adapted from Chang and Larracuente, 2019). The centromere (C), long arm (Y^L^), and short arm (Y^S^) of the chromosome are indicated. The sites assayed in E are connected to their location within a scaffold or contig. (G-J) Karyotypes from larval neuroblasts stained with DAPI. Dot chromosomes are indicated by yellow carets, Y chromosomes are indicated by teal carets, and 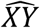 chromosomes are indicated by magenta carets.

False positive effects are a major challenge in targeting the Y chromosome because irrelevant genetic changes to the chromosome (1) cannot be purged due to the lack of recombination, (2) are likely to influence male fertility due to the prevalence of loci that influence fertility on the Y chromosome, and (3) can accumulate unchecked in the compound chromosome background where there is less selective pressure (Cook *et al*. 2010b). To compensate for this we screened through a large number of flies to obtain multiple independent alleles of each gene, thereby increasing confidence in any observed phenotype. Once mutants were obtained the edited Y chromosome was removed from the compound background (Fig. 1D). Furthermore, any quantitative fertility tests were completed on at least three mutant alleles and compared with three wild-type alleles.

### PRY mutants show reduced fertility but are not sterile

*PRY* is Y-linked in both *Sophophora* and *Drosophila* groups and therefore moved onto the Y chromosome, where it has been evolving, over 50 million years ago (Koerich *et al*. 2008). This long history of male-limited inheritance and testis-biased expression would suggest a role in male fertility. Yet V24, a fertile translocation line, has a breakpoint in the *PRY* gene (Carvalho *et al*. 2000), suggesting that the *PRY* gene is not required for male fertility.

To more directly test whether *PRY* plays a quantitative role in male fertility we used CRISPR to create *PRY* mutants. We targeted 237 bp of exon 2 of *PRY*, near the beginning of the PKD/REJ Domain 1, the first conserved domain in the protein (Fig. S1A). We used the BDC line described above to avoid biasing against mutants that might be less fertile than the control. We were able to recover five alleles of *PRY* including four that were derived from independent F0 flies and four that were frameshifts (Fig. S1C, Table 2). RT-PCR of *PRY G54.1* using primers downstream of the guide sites indicates that the RNA transcript is still present despite the frameshift (Fig. S4A,C). We can therefore rule out mechanisms such as nonsense mediated mRNA decay. We nevertheless expect that deletion or scrambling of ∼82% of the coding sequence would result in loss of function. We then removed the compound chromosome background and checked whether XY males carrying *PRY* mutations were sterile. Most of the males tested produced offspring, so we concluded that *PRY* was not required for male fertility. This is consistent with the fertility seen for the V24 translocation line.

We next chose what we expected to be the three strongest *PRY* alleles – those with the largest deletion and frameshifts – to test for more subtle effects of the *PRY* gene on male fertility. To control the genetic background, we first outcrossed the mutant alleles to Canton-S for a total of 5 generations (Fig. 1D). Because we selected males at each generation, there is no recombination and an 87.5% chance of establishing an entirely wild-type Canton-S background (note: the Y chromosome also originated from Canton-S). For controls, we concurrently outcrossed males from the original BDC stock, and established three independent control lines. Because mutations accumulate rapidly on redundant Y chromosomes (Cook *et al*. 2010a), we did not retain the mutants in the compound background.

To measure fertility, fecundity and hatchability more quantitatively, we crossed single Canton-S females to mutant or control males. We removed the males after a single mating and maintained the isolated females for five days. We counted the number of eggs laid each day (fecundity) and the number of pupae formed from those eggs one week later (fertility). Results are summarized by allele and replicate in Table S4. Because the Y chromosome contains so many loci that affect fertility and does not undergo recombination, the fertility effects of a specific mutation can easily be obscured by Y-linked off-target effects from CRISPR or preexisting variation. To rule out such effects we first compared the egg and pupal counts of the independently derived alleles of each genotype by day (Fig. S5A,C,E) and cumulatively (Fig. S6A,C). To account for the possibility of Y-linked fertility effects unrelated to *PRY*, we statistically confirmed that the counts were not statistically different among alleles of the same genotype using a Kruskal-Wallis Rank Sum test (Table S6). We then statistically tested for differences between the genotypes – control and *PRY* – using a Wilcoxon rank sum test (Table S7, Fig. 2A,C,E).

A small yet consistent defect was seen in the fecundity of *PRY* mutants that peaked on Day 3 after mating, resulting in significant differences in the cumulative counts. A strong and significant defect was seen in the fertility of *PRY* mutants on all five days and in the cumulative counts. The defect in egg and pupal counts could be caused by a defect in sperm production levels, ability to transfer sperm to females, ability of sperm to enter storage in the female, or sperm motility (and thereby fertilization ability) in mutants. Follow-up studies will be necessary to further dissect the specific role of *PRY* in male fertility.

### FDY and vig2 do not appear essential to regulate male fertility

As the youngest gene on the Y chromosome, *FDY* presents an opportunity for understanding the characteristics that allow a gene to survive on the Y chromosome and for characterizing early changes that occur when a gene adapts to the Y chromosome environment. *FDY* is part of an 11 kb segment that is syntenic and similar in sequence to a segment on the third chromosome. This segment is found on the Y chromosome in *Drosophila melanogaster* but none of the closely related species, suggesting a duplication event occurred in the *melanogaster* lineage the past 2.5 million years (Carvalho, 2015). *FDY* is the only gene that shows evidence of selection in this segment (Carvalho, 2015), and most of the other protein-coding genes have degenerated. *FDY* has a 96.5% amino acid sequence identity to its third chromosome paralog, *vig2*, which presented several challenges for targeting the gene.

We designed sgRNAs to target the 5’ end of the gene, including the transcription start site (Fig. S2). Due to the high level of sequence similarity, our sgRNAs target both *FDY* and *vig2*, but we were careful during outcrossing to recover only wild-type alleles at *vig2*. By design, our sgRNAs should delete the transcription start site of *FDY* but not *vig2*. We again used the compound chromosome strategy to avoid biasing against mutants that might be less fertile.

We isolated several independent alleles of *FDY*. Six of the independently-derived mutants had an identical deletion and upon close inspection of the region we noticed an eight base pair microhomology domain at each of the two sgRNA sites. The common deletion in the six independent lines resulted from repair through binding of these homologous regions (Fig. S2). In total we created three distinct independently-derived alleles. All of the alleles eliminate the transcription start site and two disrupt the reading frame (Table 2). While we cannot be sure that these mutants are null, at least one-third of the coding sequence, including half of a conserved Hyaluronin/mRNA-binding protein domain are deleted in our mutants (Fig. S2). We expect such a perturbation to have strong effects on FDY protein function.

*FDY* is not in one of the regions of the Y previously shown to be required for male fertility, but might have a measurable role in male fertility as *PRY* did. We crossed our CRISPR mutants to Canton-S to isolate the Y chromosome from the compound chromosome background (Fig. 1D). As expected, none of the alleles were sterile. We wondered whether there may be some functional redundancy between *vig2* and *FDY. vig2* has ubiquitous expression, including in the testis, but no reported role in male fertility. The *vig2*^*PL470*^ mutant contains a PiggyBac insertion in the *vig2* gene that eliminates detectable protein (Gracheva *et al*. 2009) and transcript (Fig. S4D,E) expression but is viable and not sterile (Gracheva *et al*. 2009). We found that *FDY*-*vig2* double mutants were also viable and not sterile (Table S9).

To test for more subtle fertility effects of *FDY* we measured fertility, fecundity and hatchability of females mated to *FDY* males and corresponding control lines (Table S5), as was done for *PRY*. The Kruskal-Wallis Rank Sum test indicated a significant difference in the fecundity and fertility of a single control allele, E. There was little variation observed between control alleles, A and C, or the *FDY* mutant alleles (Fig. S5B,D,F). The control alleles were derived from flies that were injected with guide RNAs for *FDY* but were negative for mutations in both the *FDY* and *vig2* locus, and therefore could contain off-target mutations, other CRISPR side-effects (see below), or variants that are Y-linked. Regardless of its cause, the increased fertility of the outlier allele (1) was not removed during out-crossing (Fig. 1D) and (2) is not related to *FDY*. Table S7 includes the results of the Wilcoxon Rank Sum Tests to compare control and *FDY*, both with and without the outlier. However, we base our subsequent conclusions and discussion on removal of this outlier allele. Per-day fertility, fecundity and hatchability are plotted in Fig. 2B,D,F and cumulative results are plotted in Fig. S6B,D showing no significant differences (Table S7) in egg or pupal counts between females mated to *FDY* or control males. *FDY* mutant males also had no obvious effects on the variability of counts of eggs laid or pupae produced (Table S8). *FDY* may yet have effects on fertility that are too subtle to detect or limited to stressful or competitive contexts.

### Y chromosome truncation is an artefact of FDY targeting by CRISPR

After injection with *FDY* sgRNAs, roughly 20% of F1 flies appeared to have a large truncation of the Y chromosome that did not correspond to the intended deletion. We were able to visually detect the resulting significant change in heterochromatin levels through an eye color marker – *white*^*mottled4*^ (*w*^*m4*^) – that was present on the compound 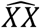 in our BDC line. The marker is intended to distinguish between females that carry a Y chromosome (suppressed variegation – Fig. 3B) from those that do not (strong variegation – Fig. 3A). Strong variegation, likely associated with occasional Y chromosome loss, was observed very rarely in the parental BDC line. However, with *FDY-*targeting we observed a large number of flies with an intermediate level of variegation (Fig. 3C,D), which was not observed in any other line. The intermediate variegation was stably heritable and tracked with the Y chromosome (see Fig. 1B,C). Therefore, it could not be caused by mutation of *vig2*. Three further lines of evidence suggested that the change in variegation was caused by truncation of the Y chromosome.

First, a PCR survey (Fig. 3E,F) indicated that *FDY* and other loci on the long but not short arm of the Y chromosome were absent in intermediate-variegating 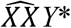 females. Two loci on the short arm were present in all of the females tested, and two loci on the long arm were absent from all of the females tested (Fig. 3F). Most variegating lines successfully amplified the 3’ end of *FDY* – suggesting that this region was intact. All of these variegating lines failed to amplify the target site. This suggests a deletion of most of the long arm (Fig. 3F), ending near the target site within *FDY*.

Second, Y chromosomes from flies with intermediate variegation appeared smaller in karyotypes (Fig. 3G-J). To easily distinguish the maternal Y chromosome from the paternal Y chromosome we crossed the variegated 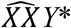 to an 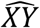 male. We squashed brains of male larvae from this cross and examined condensed chromosomes in neuroblasts. In normal 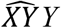 males we observed a Y chromosome that was approximately equal in length to the autosomes and had characteristic condensed regions (Fig. 3G,H). In contrast, 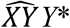 males had all of the other chromosomes but no characteristic Y chromosome (Fig. 3I,J). Instead they had an extra small chromosome body, approximately the length of the dot chromosome. This truncated chromosome was seen in the karyotypes of all three independently-derived lines with intermediate variegation that we examined.

Finally, males with the truncated Y chromosome were sterile when crossed out of the compound chromosome background. This is consistent with the loss of four fertility factors as predicted by the PCR survey. Our results could be explained by either improper repair of double-strand breaks induced by the sgRNAs at the *FDY* locus or an exchange between cut sites in *FDY* and *vig2*. In either case, DNA segments attached to the centromere would be expected to be retained while those distal from the break site would be expected to be lost. This peculiar artefact thus allows us to infer that *FDY* is located on the long arm of the Y chromosome. Furthermore, it demonstrates that the BDC background is a viable strategy for maintaining recessive sterile Y-linked mutations.

### CCY CRISPR mutants result in male sterility even in the presence of a wild-type copy of CCY

*WDY, ORY* and *CCY* were genes identified by genome sequencing and mapped by PCR to the *kl-1, ks-1*, and *ks-2* fertility factor intervals, respectively (Carvalho *et al*. 2001; Vibranovski *et al*. 2008, Table S1). While these are currently the only likely candidates in their respective interval, less than ∼50% of the sequence of the Y chromosome is currently known. Because the fertility factor intervals are very large, e.g. ∼1.2 Mb for *ks-2* (Kennison 1983), it remains possible that an undiscovered gene in the region is responsible for the sterility phenotype.

We used our CRISPR strategy to create *CCY* mutants in order to test whether *CCY* is required for male fertility. We used sgRNAs 3 and 4 to excise 554 bp of exon 1 (Fig. S3), removing a large portion of a conserved region with homology to the SbcC, Coiled-coils, and MIT-CorA domains. We observed a high level of sterility in the F0 generation (see Fig. 1B,C for crossing scheme) that could perhaps be dismissed as these flies were injected as embryos. Surprisingly, however, we observed strong male-biased sterility in the F1 generation (see Fig. 1C). 64.6% (82/127) of F1 males inheriting the edited Y were sterile while only 6.7% (2/30) females were sterile. The level of female sterility is not unreasonable for single fly crosses, but the level of sterility in males is very striking. We did not notice significant sterility in F1 males when targeting any of the other Y chromosome genes. The male sterility in the F1 generation is a dominant effect as all of the F1 males have one wild-type copy of the Y from the 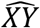 (see Fig. 1C). As a result of this dominant sterility we were not able to establish or maintain lines of any *CCY* mutants. Although F1 females with the *CCY* mutant Y chromosomes were fertile, the *CCY* mutant Y chromosomes were only recovered in their sons (see Fig. 1B), which were sterile, and not in their daughters. In the F0 generation (see Fig. 1B,C for crossing scheme) band-shifts were observed, indicating deletion at both sgRNA sites. Although no band shifts were observed in F1 or later generations of both sterile and non-sterile F1 flies, sequencing the region revealed the presence of several small insertion-deletions (indels) at one of the sgRNA sites. All of the indels we identified that changed the coding sequence by a multiple of 3 (thereby preserving the coding frame) resulted in non-sterile flies. In contrast, all of the indels we identified that were not a multiple of 3 (thereby disrupting the coding frame) resulted in sterile flies (Table S10, Table 3). This may indicate that an intact reading frame for *CCY* is necessary for male fertility.

We found no evidence of Cas9 cuts at sgRNA 4 past the F0 generation. To rule out off-target effects specific to this sgRNA, we used a different 3’ sgRNA – sgRNA 1, in another attempt at making a *CCY* mutant. We again observed high levels of dominant male sterility. Although we were able to retain one line with a 180 bp deletion, the deletion is in-frame, does not disrupt the Coiled-coils or MIT-CorA domain, and therefore is unlikely to disrupt the protein’s function in a meaningful way. This single allele had no detectable effects on male fertility (data not shown) and was not retained.

### Knockdown of the predicted fertility factors in the testis causes sterility

As an alternate method to test the requirement of the predicted fertility factors in male fertility, we crossed publicly available RNAi lines targeting each factor to Bam-Gal4, a germline-specific driver (Chen and McKearin, 2003). A *UAS-Dicer2* was present with the driver and presumably boosted knockdown efficiency. We then tested the knockdown males for sterility. Knockdown of all six predicted fertility factors resulted in sterile males, including *kl-3*, and *kl-5*, as expected, and *kl-2, ORY, WDY* and *CCY*, as expected (Table 4). In particular three independent RNAi constructs for *WDY* confirmed its requirement for male fertility. In contrast, knockdown of *FDY* and *PRY*, which our CRISPR data showed to not be required for fertility, did not result in sterile males (Table 4). We used RT-PCR to measure transcript levels in adult males after *PRY* knockdown (Fig. S4B,C). Although some transcript is detectable in the *PRY* RNAi line, more than 75% of the transcript is knocked down. We were unable to test the level of knockdown of *FDY*, however, the absence of a detectable phenotype of *FDY* RNAi is consistent with that of the CRISPR mutant. In summary, our RNAi results support the current assignment of fertility factor genes.

The sensitivity of male fertility to the expression of fertility factor genes is striking. RNAi lines often vary widely in the degree of knockdown they produce. However, each of the 8 RNAi lines we tested caused 100% penetrance of male sterility (Table 4). Either these RNAi lines all produce highly efficient knockdown by chance, or there is a strong dosage sensitivity for these genes.

### The role of CCY in spermatogenesis

Hardy and colleagues investigated defects of males with segmental deletion of *ks-2* (Hardy *et al*. 1981) and found that they fail during sperm individualization. We tested whether knockdown of *CCY*, the gene predicted to be responsible for the sterility of *ks-2*, also disrupted spermiogenesis at the individualization stage. We expressed *CCY* RNAi in the testis with *Bam*-Gal4. Semi-quantitative RT-PCR demonstrated that *CCY* gene expression is reduced by more than 90% in the testes of these knockdown males (Fig. S4F,G).

Individualization involves an initial stage in which an actin-rich individualization cone forms around spermatid heads, a middle stage where the cone begins to progress, and a final stage in which cytoplasmic material and the individualization cone accumulate at the base of the of spermatid cluster in a “waste bag” structure that is degraded (Fabrizio *et al*. 1998; Rathke *et al*. 2007; Fabian and Brill 2012). Males knocked down for *CCY* had no mature sperm in their seminal vesicles (Fig. 4J,K). In the knockdown testis we never saw cystic bulges (identified as actin-staining individualization cones with no immediately adjacent nuclei) or wastebags (Fig. 4I), which characterize spermatids in late and post-individualization stages. We did, however, observe protamine accumulation (Fig. 4A-D), initial individualization cone formation (Fig. 4E,F), and early progression of the individualization cone (Fig. 4G,H). Upon close examination of this early progression of the individualization cone, we noticed disorganization of the spermatid bundles relative to wild type (arrowheads, Fig. 4F,H). We therefore conclude that *CCY* knockdown spermatids fail during the progression of the individualization cone.

**Figure 4.**
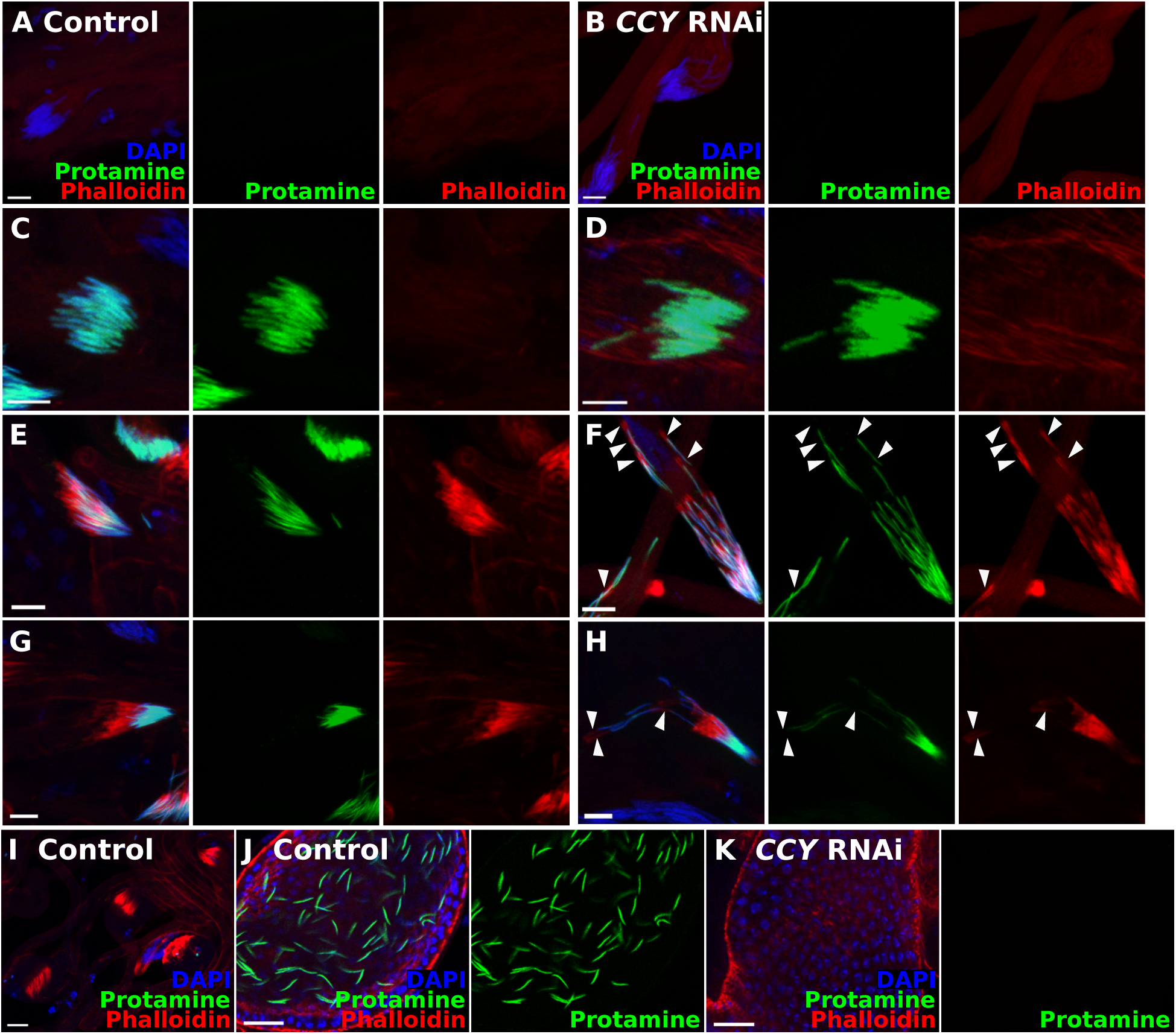
RNAi of *CCY* disrupts spermiogenesis at the individualization stage. Testis squashes from flies containing the Bam-Gal4 driver (Chen and McKearin 2003) and either *attP40* (Control) or *UAS-CCY* RNAi. A protamine-GFP reporter is present (mature sperm heads in green) (Manier *et al*. 2010) and the preps have been stained with DAPI (nuclei in blue) and phalloidin (Actin in red). Stages of spermatogenesis: (A,B) early canoe stage (protamine negative, Actin negative) (C,D) late canoe stage (protamine positive, Actin negative) (E,F) individualization cone formation (protamine positive, Actin positive) (G,H) individualization cone progression (Actin displaced from nuclei) (I) waste bags (J,K) mature sperm in seminal vesicle. Scale bars : A-H 10µm, I-K 20µm

## DISCUSSION

Here we report fertility phenotypes of CRISPR mutants in three Y chromosome genes: *PRY, FDY*, and *CCY*. The mutants were made using CRISPR together with an effective genetic strategy for obtaining Y-linked mutants that are sterile or semi-fertile. The fertility phenotype of *PRY* shows that established genes on the Y chromosome, beyond fertility factor genes, can contribute to male fertility. Furthermore, the dominant sterility following targeting of *CCY* and the sterility of RNAi of all predicted fertility factors illustrates the critical importance of these factors in spermatogenesis. Complications in constructing the mutants have been instructive – raising important questions about our current understanding of the genetics of Y chromosome genes.

### PRY activity contributes to male fertility while FDY activity does not appear to

All Y-linked genes have testis-biased expression and male-limited inheritance, and therefore Y-linked genes may be expected to influence male fertility. Yet only six Y-linked genes are thought to be required for male fertility. *PRY*’s location at the breakpoint of a fertile translocation line suggests that it is not required for male fertility. We found that both egg-laying and the number of pupae produced by females mated to *PRY* mutant males are significantly reduced relative to control, although the mutant males are still fertile. An egg-laying defect is surprising given that egg-laying is regulated by seminal fluid proteins from the accessory glands (Avila *et al*. 2011). Yet *PRY* has domain homology to polycystins which have been linked to sperm storage and fertilization (Kierszenbaum 2004; Köttgen *et al*. 2011; Yang and Lu 2011). Reduced sperm in the female reproductive tract would result in reduced levels of the sperm-binding protein, Sex Peptide, which stimulates egg production and egg laying (Chapman *et al*. 2003; Liu and Kubli 2003). Identifying whether *PRY* defects occur in sperm production, transfer, motility, or storage and how *PRY* regulates such processes will be the next step in understanding the function of this gene.

We do not see a fertility defect in *FDY* mutants, though we have not tested the mutants in competitive assays, such as sperm competition situations, or under stressful conditions, such as extreme temperatures. The close homology to *vig2* might suggest a role for *FDY* in chromatin regulation since *vig2* is (1) a modifier of position effect variegation (2) regulates levels of H3K9me2, a marker of constitutive heterochromatin (Gracheva *et al*. 2009; Schneiderman *et al*. 2010), and (3) Vig2 protein interacts with heterochromatin protein 1 (HP1) and the histone cluster (Tsui *et al*. 2018). *FDY* has a more limited tissue expression pattern than *vig2*. Testing whether *FDY* is involved in chromatin regulation in cells where it is expressed will be an important future direction as it may reveal a function for a Y-linked gene beyond fertility. Statistical tests showed a significant difference in fertility and fecundity of one of the *FDY* control alleles. This could be due to a rare variant in the parental stock that is represented in this allele but not in any of the other control or *FDY* alleles. Alternatively, the outlier could be due to an off-target or side-effect of CRISPR. Regardless of its source, the variation observed in the single control allele is not present in the other two control alleles or in the three mutant alleles and therefore cannot be related to *FDY*. We mention this outlier for the sake of transparency and to highlight an important consideration for future studies – the importance of testing multiple alleles of each genotype for phenotypic analysis of the Y chromosome in order to rule out false positive effects on fertility.

### Fertility factor genes and function

Our RNAi data suggest that all 6 genes previously proposed to be fertility factors (Carvalho *et al*. 2000, 2001; Vibranovski *et al*. 2008) were correctly assigned. Furthermore, RNAi suggests that the amount of product of these genes is relevant to their fertility function, rather than a structural feature, such as a Y chromosome loop or intronic repetitive element. Three of the fertility factors encode dyneins which are thought to be structural proteins of the sperm flagellum required for sperm motility (Goldstein *et al*. 1982; Gepner and Hays 1993), but the mechanism of the remaining fertility factors is still unknown. Hardy *et al*. (1981) showed that segmental deletion of several fertility factors led to defects in primary spermatocytes and spermatid failure at individualization (Hardy *et al*. 1981). We show, more specifically, that in *CCY* RNAi lines spermatids fail during progression of the individualization complex. This may indicate that CCY protein, either directly or indirectly, may affect cytoskeletal dynamics (Steinhauer 2015).

### CCY-targeting leads to dominant sterility

One of our most surprising findings is that targeting *CCY* leads to dominant male sterility. Dominant sterility is rare, but is more common on the Y chromosome (Lindsley and Tokuyasu 1980). The dominant sterility may be caused in several ways: (1) an off-site translocation, (2) loss of *CCY* function, (3) the formation of an aberrant gain-of-function product, or (4) a genetic interaction.

Kennison reported 50 Y-linked dominant male-sterile alleles after EMS and X-ray mutagenesis (Kennison 1983). He argued that the dominant sterility was the result of Y:autosome translocations, similar to the dominant sterility caused by X:autosome translocations in Lifschytz and Lindsley (1972). Consistent with this idea, all 50 dominant alleles he examined showed pseudolinkage with a visible autosomal marker (Kennison 1983). He observed these dominant sterile translocations with breakpoints on both the long and short arm of the Y. We can rule out a translocation breakpoint in the target region of *CCY* in at least a subset of the sterile 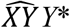 mutants: double peaks indicate amplification and intact sequences in both the single Y chromosome and the compound Y chromosome. However, we cannot rule out translocations in other Y-linked regions of these mutants or in other mutants. The idea that Y:autosome translocations may cause defects in spermatogenesis is interesting and has not been investigated. Chang and Larracuente recently documented duplications of exons of several Y chromosome genes (Chang and Larracuente 2019). While they did not find duplications of *CCY* exons, such duplications may exist in the unsequenced portions of the genome. CRISPR editing of a duplicated region may result in a chromosome rearrangement that could dominantly disrupt spermatogenesis.

Loss of *CCY* function may also cause dominant sterility due to extreme dosage sensitivity for the gene. The high penetrance of sterility in the RNAi lines supports the idea of dosage sensitivity for all the fertility factors. The absence of intended deletions in mutants after the F0 generation may also indicate a sensitivity of the locus to perturbation. In this case, the sterility observed in the presence of a normal Y chromosome would have to indicate that the levels of *CCY* are calibrated to the number of Y chromosomes present. In other words, one wild-type copy of *CCY* is necessary per Y chromosome present.

Another explanation for the dominant sterility is the creation of an aberrant “dominant negative” or antimorphic product. Dominant sterility was seen with frame shifts of both +1 and +2 base pairs, making it unlikely that an aberrant product arises from an alternative reading frame. Rather, it may arise from an alternative start site of the main coding frame resulting in a truncated RNA or protein product that may be toxic. Antibodies to the C-terminus of the protein may be able to detect such a product. We do not favor this explanation as antimorphic mutations are generally considered to be rare, whereas Kennison observed dominant sterility at a high frequency and on both arms of the Y chromosome (Kennison 1983).

Finally, apparent dominant sterility could be caused by a genetic interaction, such as between a lesion in *CCY* and a rearrangement associated with the 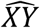 chromosome. Genetic interactions between a Y-duplication and X-deletion (Rahman and Lindsley 1981) and between a Y-autosome translocation and a X-deletion (Lindsley and Tokuyasu 1980) were previously shown to cause sterility.

An entirely different strategy would be required to maintain lesions that result in dominant sterility in order to study their mechanism. Because we used a compound 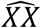 chromosome in our strategy, there was no way to maintain the Y chromosome in females. A strategy like that of Kennison (1983), that allows the Y chromosome to be maintained in the female line is necessary. Although we were not successful in maintaining a CRISPR mutant of *CCY*, our data provide strong evidence that an intact *CCY* coding frame is required for male fertility.

### Truncation at the FDY locus

Large Y chromosome truncations appear to form regularly upon double-strand breaks at the *FDY* locus. This may indicate a deficit in double-strand break repair at the *FDY* locus or could be caused by translocation between *FDY* and *vig2*. Because of the extremely high sequence similarity between these genes, our sgRNAs are expected to cut at both sites. The expected translocation would replace most of the long arm of the Y chromosome with a small segment of the third chromosome. The reciprocal translocation would replace most of 3R with the long arm of the Y chromosome and would be expected to be rapidly lost. If this explanation is correct, it may indicate a method for reproducibly creating specific translocations. Indeed Lynagh and colleagues showed that Cas9-induced breaks in non-homologous chromosomes can induce translocations in Arabidopsis (Lynagh *et al*. 2018). Aside from the mechanism of formation of these truncations, the consistent pattern of loss of several long-arm Y chromosome sites in the truncations suggests that *FDY* is located on the long arm of the Y chromosome proximal to these sites. Importantly, our ability to maintain these truncations in our BDC background demonstrates that our strategy for retaining recessive sterile Y-linked mutations works reliably.

## CONCLUSIONS

Our work demonstrates that the Y chromosome harbors important fertility regulators beyond the six previously identified fertility factors, expanding our understanding of the role of the Y chromosome in regulating male fertility. We devised a strategy for creating and retaining recessive sterile mutations on the Y chromosome. However, we encountered several challenges in using this strategy to target specific genes, including dominant sterility and truncation of the Y chromosome. These challenges are indicative of the poor current understanding of the biology of this gene-poor, repeat-rich, and highly heterochromatinized chromosome. Despite its mysterious biology, the Y chromosome is not just a genetic anomaly. It plays a vital role in male fertility, adaptation, speciation and genomic conflict and is therefore an important key to better understanding evolutionary processes.

## ACKNOWLEDGEMENTS

We thank M. Zhang, A. Wong, J. Champer, A. Jain, G. Chi, T. Nguyen, I. Amaro, and N. Buehner as well as the members of the Clark and Wolfner labs for experimental assistance and/or advice. We thank S. Delbare, M. Munasinghe, S. Misra, Y. Ahmed-Braimah, and three anonymous reviewers for discussion and comments on the manuscript. Stocks obtained from the Bloomington Drosophila Stock Center (NIH P40OD018537) were used in this study. This work was supported by funds from the Cornell Civil and Environmental Engineering Department and a grant from the National Institutes of Health (R01 GM119125 to A. Clark and D. Barbash).

